# Spindle-locked ripples mediate memory reactivation during human NREM sleep

**DOI:** 10.1101/2023.01.27.525854

**Authors:** Thomas Schreiner, Benjamin J. Griffiths, Merve Kutlu, Christian Vollmar, Elisabeth Kaufmann, Stefanie Quach, Jan Remi, Soheyl Noachtar, Tobias Staudigl

## Abstract

Memory consolidation relies on the reactivation of previous experiences during sleep. The precise interplay of sleep-related oscillations (slow oscillations, spindles and ripples) is thought to coordinate the information flow between relevant brain areas, with ripples mediating memory reactivation. However, in humans empirical evidence for a role of ripples in memory reactivation is lacking. Here, we investigated the relevance of sleep oscillations and specifically ripples for memory reactivation during human sleep using targeted memory reactivation (TMR). Intracranial electrophysiology in epilepsy patients and scalp EEG in healthy participants revealed that elevated levels of SO-spindle activity promoted the read-out of TMR induced memory reactivation. Importantly, spindle-locked ripples recorded intracranially from the medial temporal lobe were found to be instrumental for memory reactivation to unfold during non-rapid eye movement (NREM) sleep. Our findings establish ripples as key-oscillation in human systems consolidation and emphasize the importance of the coordinated interplay of the cardinal sleep oscillations.

## Introduction

Contemporary models propose that memory consolidation, i.e., the strengthening of memories during sleep, is achieved by reactivating experiences that were encoded during prior wakefulness ^1,2^. Through reactivation, memories are relayed between the hippocampus and cortical long-term stores, transforming initially labile memories into long-lasting ones ^3^. The essential communication between the hippocampus, thalamus and cortex, as well as the strengthening of memories in cortical networks, is thought to be facilitated by a precise temporal coordination between the cardinal non-rapid eye movement (NREM) sleep related oscillations, namely cortical slow oscillations (SOs), thalamocortical sleep spindles and hippocampal ripples ^4–6^.

SOs (< 1 Hz), initiate time windows of excitability and inhibition not only in cortical but also in subcortical areas ^7–9^. They ignite the generation of sleep spindles in the thalamus, which nest in the excitable upstates of cortical SOs ^10,11^. Spindles (12 – 16 Hz), in turn, have been shown to gate Ca2+ influx into dendrites, putatively facilitating synaptic plasticity in cortical areas ^12–14^. Lastly, hippocampal sharp-wave ripples (80 – 120 Hz in humans) are assumed to coordinate neural population dynamics in the hippocampus to reactivate recently formed memories ^15,16^. Ripples tend to occur during the excitable troughs of spindles^17,18^. The formation of such spindle-ripple events is thought to facilitate the transfer of reactivated memories to the cortex ^19,20^. Hence, while SO-spindle coupling is assumed to ensure that cortical target areas are optimally tuned for synaptic plasticity when memories are reactivated, memory consolidation ultimately relies on ripples to trigger and coordinate memory reactivation processes both in the hippocampus and cortical long-term stores ^16^.

Studies using intracranial recordings in epileptic patients have established the hierarchical synchronization of SOs, spindles and ripples during human NREM sleep ^17,21–26^. However, whether spindle-locked ripples contribute to memory consolidation by mediating memory reactivation in humans is currently unknown. Here, we set out to assess the relevance of sleep oscillations and specifically sharp wave ripples for memory reactivation during human NREM sleep. We recorded scalp EEG in healthy participants and intracranial EEG in epilepsy patients while they retrieved real-world spatial memories (i.e., prior learned head orientation – image associations). Importantly, head orientations were linked to specific sound cues, which were presented again during subsequent non-rapid eye movement (NREM) sleep to trigger the reactivation of head orientation-related memories (i.e., targeted memory reactivation, TMR ^27^). Using multivariate classification, we find that head orientation-related electrophysiological signatures are reactivated during successful awake memory retrieval as well as during TMR while participants were asleep. During sleep, elevated levels of SO-spindle activity promote the read-out of memory reactivation in both scalp and intracranial EEG recordings. Leveraging direct access to medial temporal lobe (MTL) electrophysiology in epilepsy patient, we show that spindle-locked ripples are instrumental for memory reactivation to unfold during human sleep, establishing a role of sharp wave-ripples for memory reactivation in humans.

## Results [EEG – healthy participants]

Twenty-five participants (age: 25.2 ± 0.6; 16 female) took part in the scalp EEG study. Experimental sessions started in the evening around 7 p.m. After an initial training phase (see Methods), participants performed a real-world spatial memory task, where they learned to associate 168 items (images of objects) with specific head orientations (see Fig. 1a). Importantly, a specific sound cue was assigned to each of the four non-central head orientations. After a delay filled with a distractor task, memory performance was tested in a stepwise manner. First, participants made object-recognition judgments for all old items, randomly intermixed with new items. Then, for recognized items only, participants indicated which of the four head orientations was associated with the item during the learning phase (associative retrieval, Fig. 1a). After finishing the memory task, participants went to sleep. During one hour of NREM sleep, two out of the four sounds (one sound associated with the right-sided and one with the left-sided head orientations, respectively) were repeatedly presented as TMR cues, while an additional sound, unrelated to any learning, served as a control sound. We reasoned that presenting TMR cues during sleep would ignite reactivation of the related head orientations and the associated items.

**Fig. 1.**
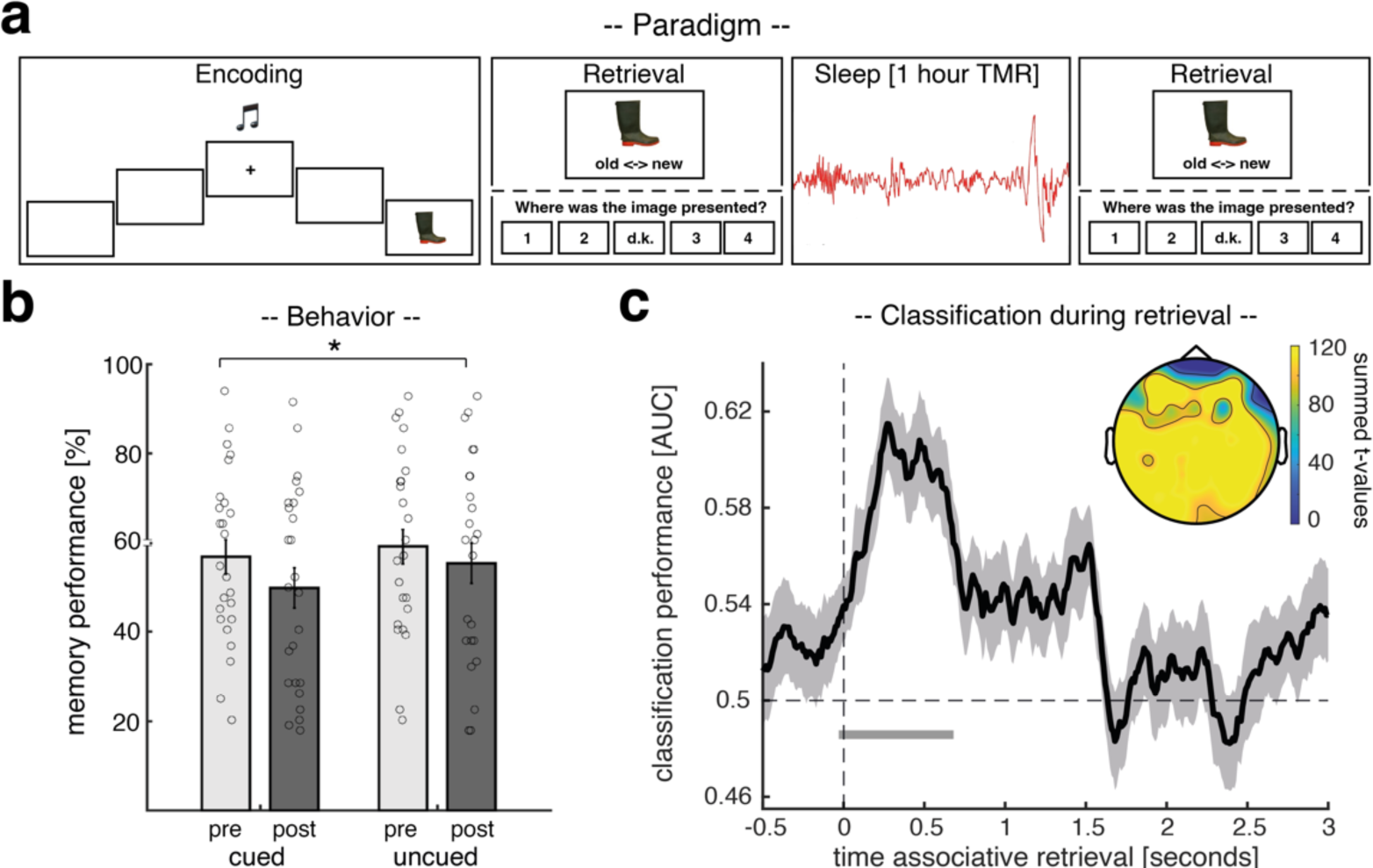
Experimental procedure, behavioral results, and retrieval locked reactivation of head orientations. (a) During encoding, participants were consecutively presented with 168 images (EEG study) / 144 images (intracranial EEG study) of objects on four flanking screens (positioned at −60°, −30°, +30° and 60° relative to the center screen). Participants turned their head towards the relevant screen, cued by one of four orientation-specific sounds. Memory performance was tested via a recognition test followed by an associative retrieval (this procedure was used before and after sleep): First, participants made object-recognition judgments (old or new). Then, for recognized images only, participants indicated which of the four head orientations was associated with the item during the learning phase. During NREM sleep, two of the learning-related sounds (one related to left-sided and one related to right-sided head orientation) and one control sound, which was not part of the learning material, were presented for 60 minutes. (b) Behavioral results for both experimental sessions pre- (light gray) and post-sleep (dark gray), separated into cued and uncued trials. Bar graphs show mean (±SEM) percentage of recalled head orientations. Dots indicate individual memory performance of participants (N = 25). The star denote the significant interaction(pre vs. post x cued vs. uncued) as derived from a repeated measures ANOVA (F1,24 = 5.48; *p* = 0.028). (c) Later cued head orientations (left vs. right) could be reliably decoded (above chance) from the retrieval data, starting around the onset of the associate prompt (the black solid line indicates decoding performance (±SEM)). The horizontal dashed line indicates chance level performance (i.e., 0.5). The vertical solid line indicates the onset of associative retrieval trials (time = 0). The lower horizontal gray line shows the temporal extent of significant decoding results as derived from a dependent-samples t-test (two-sided, *p* < 0.001, cluster corrected across time). The topographical insert illustrates the results of a “searchlight decoding procedure”, indicating that bilateral centro-parietal and occipital areas exhibited stimulus-category related effects (please note that statistical tests were done for illustrative purposes only).

### Behavioral results

To test for potential differences in memory performance between test times and TMR conditions, we conducted an ANOVA for the cued recall, including the factors cueing (cued vs. uncued) and test-time (pre- vs. post-sleep). Results indicated that memory performance declined over the course of sleep (main factor test-time: F_1,24_ = 19.24; *p* < 0.001). Importantly though, the interaction between test-time and cueing (F_1,24_ = 5.48; *p* = 0.028) was also significant, indicating that TMR did modulate memory performance. However, TMR did not benefit memory performance as expected ^28^, but had a detrimental effect on retrieval abilities (cued pre-sleep: 57.23 ± 3.92% vs. cued post-sleep: 50.42 ± 4.56%; uncued pre-sleep: 58.76 ± 4.13% vs. uncued post-sleep: 54.90 ± 4.61%; see Fig. 1b). Follow up post-hoc t-test (relative memory performance pre- to post-sleep) also indicated that uncued items were better remembered as compared to uncued items (t_1,24_ = 2.747; *p* = 0.011). For recognition memory, we neither found a significant main effect of test time (F_1,24_ = 0.29; *p* = 0.59); nor a significant interaction between test-time and cueing (F_1,24_ = 0.08; *p* = 0.77; see Supplementary Fig. 1 for details).

### Head orientation-related activity is reactivated during successful retrieval

Next, we set out to test whether we could decode head orientation-related activity from EEG signals during retrieval, which would allow us to track corresponding reactivation processes during TMR (see below). To extract head orientation-related patterns of neuronal activity during retrieval, we pooled the data from the associative retrieval (i.e., when participants had to remember image related head orientations) across pre- and post-sleep sessions. Furthermore, we restricted the analysis to those items whose head orientations were remembered correctly and that were selected for TMR (i.e., one left sided and one right sided head orientation per participant). We performed multivariate classification (linear discriminant analysis; LDA) on these data (Fig. 1c + d). Using fivefold cross-validation (see Methods), above-chance classification accuracy emerged around the onset of the associative memory prompt (time window: −30 ms to 680 ms; peak at 270 ms; *p* < 0.001, corrected for multiple comparisons across time). The fact that decoding accuracies ramped up slightly before the onset of the memory prompt indicate that associative retrieval processes putatively started already towards the end of old / new judgements (i.e., recognition testing); see Supplementary Fig. 2 Taken together, the retrieval data allowed us to isolate brain patterns associated with the reactivation of head orientation-related activity, which we then used to guide the analysis of memory reactivation during TMR (for results concerning the classification of later uncued head orientations during retrieval see Supplementary Fig. 3).

### TMR ignites reactivation of head orientation-related activity during NREM sleep

First, we tested whether TMR induced electrophysiological activity would discriminate between learning related and control sounds. Consistent with previous findings ^29–31^, learned TMR cues, as compared to control cues, triggered a significant power increase in the SO-spindle range (i.e., an initial low frequency burst followed by a fast spindle burst; *p* <0.001, corrected for multiple comparisons across time, frequency, and space; see Fig. 2a), foreshadowing that learning-related TMR cues might have triggered relevant neuronal processing in the sleeping brain.

**Fig. 2.**
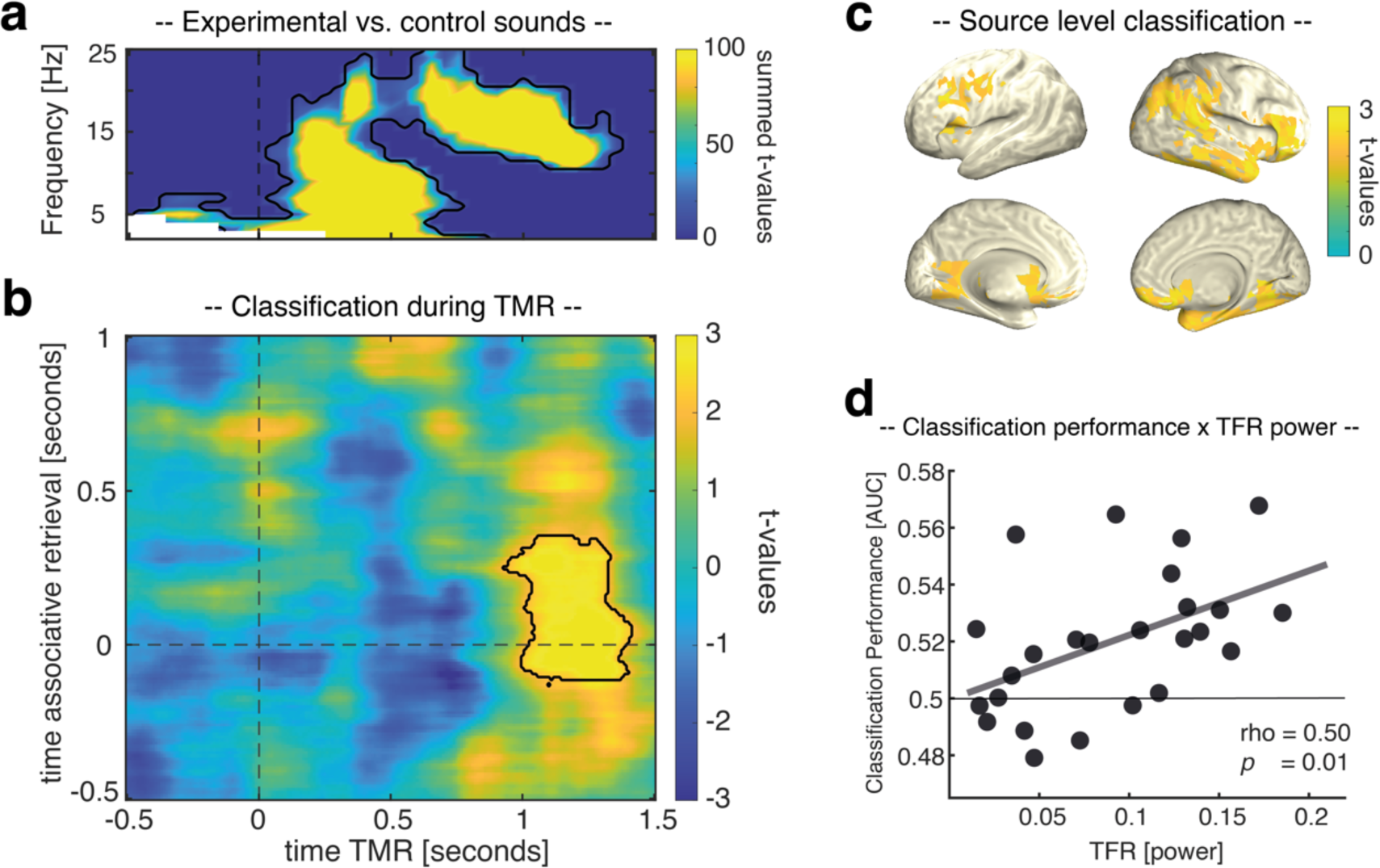
Reactivation of head orientation-related activity during TMR. (a) Power difference between learning-related TMR cues versus new control cues after statistical thresholding (*p* < 0.001, corrected) (b) Retrieval-related brain patterns (left vs. right head orientations) were decodable during TMR (contour lines indicate the extent of the cluster, *p* = 0.023 corrected; color range (blue to yellow) represents t values against chance level performance. (c) The source plots illustrate the results of a “searchlight decoding procedure”, indicating that fronto-parietal networks and the right medial temporal lobe exhibited head orientation related effects (please note that statistical tests were done for illustrative purposes only). (d) classification performance correlated positively with TMR triggered power (r = 0.50, *p* = 0.01).

To specifically test this, we next determined whether neuronal activity related to remembered head orientations would be reactivated during TMR. We first trained a classifier on the pooled associative retrieval data from both pre- and post-sleep sessions [−0.5 to 1 s]. The resulting training weights were then applied on the TMR data [-0.5 to 1.5 s]. Classifier testing labels reflected the stimulus categories used in the retrieval sessions (left- or right-sided head orientation), such that above-chance classification hallmarks TMR related activation patterns more strongly resembling the related stimulus category than the alternative stimulus category. As shown in Fig. 2b, results revealed significant above-chance classification from 930 to 1410 ms relative to TMR onset (*p* = 0.023, corrected for multiple comparisons across time), emerging during the presence of sleep spindles (associative retrieval time-window: −110 to 330 ms; the fact that decodability preceded the onset of the associative memory prompt again indicates that associative retrieval processes were probably ignited during the preceding recognition memory test). Applying the decoding procedure to source-space data revealed that these effects might have originated from fronto-parietal networks and the right medial temporal lobe (including entorhinal cortex, parahippocampus and hippocampus; see Fig. 2c). Finally, we asked whether the oscillatory fingerprint of TMR in the SO-spindle range (Fig. 2a) would be instrumental for TMR triggered memory reactivation to unfold. To address this question, we correlated, across participants, TMR triggered power (averaged across the cluster shown in Fig. 2a) and levels of mean classification performance (averaged across the cluster shown in Fig. 2b). As shown in Fig. 2d, we observed a significant positive relationship between the two variables (rho = 0.50, *p* = 0.01; for classification results based on TMR trials exhibiting increased levels of activity in the SO-spindle range see Supplementary Fig. 4).

## Results [intracranial EEG - patients]

Ten patients (age: 31.20 ± 3.46; 7 female) took part in the intracranial EEG (intracranial EEG) study. Overall, the procedures of the experiment were highly similar to the above-described scalp EEG study but optimized for patients in a clinical setting (e.g., reduced trial number in the memory task, memory task was split into three consecutive blocks; see Methods for details).

### Behavioral results

First, we tested whether the effects of TMR on memory performance, as reported above, would replicate in the patient sample. Hence, we again tested for differences in memory performance between test times and TMR conditions by conducting an ANOVA for the cued recall (factors: cueing (cued vs. uncued) and test-time (pre- vs. post-sleep)). Results revealed that patients’ memory performance also declined over the course of sleep (main factor test-time: F_1,9_ = 32.0; *p* < 0.001), comparable to the healthy participants’ decline. As in the healthy sample, we found a significant interaction between test-time and cueing (F_1,9_ = 8.28; *p* = 0.018), indicating that TMR did modulate memory performance by exerting a detrimental effect on retrieval abilities (cued pre-sleep: 58.47 ± 6.02% vs. cued post-sleep: 42.36 ± 4.89%; uncued pre-sleep: 58.88 ± 5.60% vs. uncued post-sleep: 49.58 ± 5.73%; see Fig. 3a). While the post-hoc t-test (relative change for cued vs. uncued) did not turn out to be significant (t_1,9_=1.97; *p* = 0.08), we would still like to emphasize that the overall pattern of behavioral results is highly similar to those of the healthy population. For recognition memory, there was neither a significant main effect of test time (F_1,9_ = 0.06; *p* =0.08); nor a significant interaction between test-time and cueing (F_1,9_ = 2.25; *p* = 0.16; see Supplementary Fig. 5 for details).

**Fig. 3.**
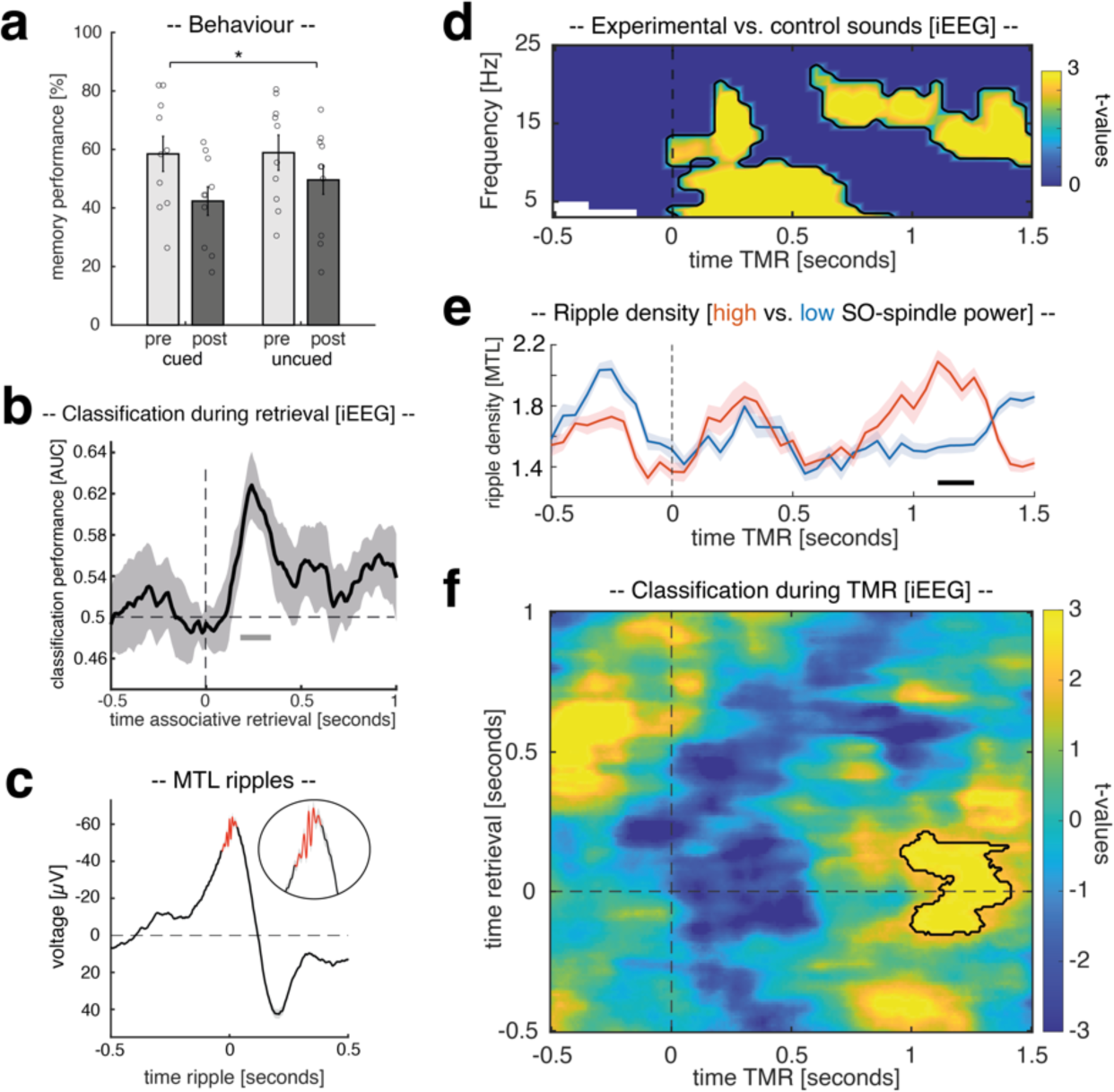
intracranial EEG study. (a) Behavioral results for both experimental sessions pre- (light gray) and post-sleep (dark gray), separated into cued and uncued trials. Bar graphs show mean (±SEM) percentage of recalled head orientations. Dots indicate individual memory performance of participants (N = 10). The star denotes the significant interaction (pre vs. post x cued vs. uncued) as derived from a repeated measures ANOVA (F1,9 = 8.28; *p* = 0.018). (b) Later cued head orientation (left vs. right) could be reliably decoded (above chance) from the retrieval data, starting around 190 ms after the onset of the associate prompt (the black solid line indicates decoding performance (±SEM)). The horizontal dashed line indicates chance level performance (i.e., 0.5). The vertical solid line indicates the onset of associative retrieval trials (time = 0). The lower horizontal gray line shows the temporal extent of significant decoding results as derived from a dependent-samples t-test (two-sided, *p* = 0.019, cluster corrected across time). (c) Ripple-triggered grand average over all detected ripples (7 patients, 14 contacts; locked to maximal negative amplitude) during TMR (-.5 to 1.5 seconds; 138.78 ± 21.72 ripples in 231.14 ± 19.94 trials). A zoomed version of the ripples is illustrated in the lower inset. The right inset shows the power spectral density (PSD) averaged across all detected SWRs [± 300 ms] indicating distinct peaks in the SO/delta, spindle and ripple range (i.e., 3 Hz, 14 Hz and 84 Hz). (d) Power difference indicate that retrieval-related TMR cues triggered increased power in intracranial EEG recordings (*p* < 0.05, corrected) as compared to control cues. (e) Ripple density for trials exhibiting high (red) and low power (blue) in the SO-spindle range, respectively. Ripple density differed significantly between conditions (*p* = 0.023; corrected), with MTL ripples peaking during elevated spindle activity. (f) Head orientation-related brain patterns (left vs. right) were decodable during TMR when contrasting high and low SO-spindle activity trials (contour lines indicate the extent of the cluster, *p* = 0.019 corrected; color range (blue to yellow) represents t values against chance level performance.

### iEEG confirms reactivation of head orientation-related activity during successful retrieval

Next, we assessed whether the intracranial data would reveal evidence for the reactivation of head orientation-related activity during retrieval, similar to the results of the scalp EEG study (see Supplementary Fig. 6 for electrode coverage of intracranial EEG recordings). Again, the associative retrieval data was pooled across pre- and post-sleep sessions, and multivariate classification (LDA) was restricted to correctly remembered items whose associated head orientations were cued during sleep (i.e., one left sided and one right sided head orientation per patient). Using fivefold cross-validation (see Methods), significant above-chance classification accuracy emerged after the onset of the associative retrieval prompt (peak at 250 ms; *p* = 0.019, corrected for multiple comparisons across time, see Fig. 3b). Hence, similar to scalp EEG recordings, multivariate classification during retrieval using intracranial EEG activity allowed us to isolate brain patterns associated with the reactivation of head orientation-related activity.

### TMR trigged reactivation of head orientation-related activity is accompanied by elevated levels of SO-spindle and ripple activity

In a first step, we tested whether TMR triggered power would also distinguish between learning related and control sounds using intracranial EEG recordings (based on frontal, parietal and temporal contacts). In line with the results of the scalp EEG study, learned TMR cues, as compared to control cues, elicited a significant power increase in in the SO-spindle range (low frequency cluster: *p* < 0.001; spindle cluster: *p* < 0.001; corrected for multiple comparisons across time and frequency, Fig. 3d).

SO-spindles have long been implicated in coordinating the emergence of hippocampal ripples and hippocampal–cortical interactions ^17,21,22,24,32^. Hence, we next tested whether different levels of cortical SO-spindle activity would influence the emergence of ripples in the medial temporal lobe (MTL). First, ripples were extracted (7 patients, 14 contacts) based on established criteria ^18^ (see Methods for details; see Fig. 3c). Then, to investigate whether activity in the SO-spindle range would affect the emergence of ripples, we sorted TMR trials as a function of power in the TFR related SO-spindle cluster (Fig. 3e) and divided the trials using a median split (see Supplementary Fig. 7 for TFR differences between high and low SO-spindle activity trials). Next, we created peri-event histograms (bin size = 50 ms) of ripple events time-locked to TMR cues for trials exhibiting high and low activity in the SO-spindle range, respectively. As shown in Fig. 3e, ripple density differed significantly between conditions (*p* = 0.023; corrected for multiple comparisons across time), with MTL ripples specifically peaking during elevated spindle activity (i.e., 1100 – 1250 ms after reminder cue onset; also see Supplementary Fig. 8). However, overall ripple number did not differ between high and low SO-spindle activity trials (high SO-spindle trials: 66.57 ± 10.57, low SO-spindle trials: 70.35 ± 10.64, t_(13)_= −1.1, *p* = 0.28), indicating that SO-spindle activity coordinates the temporal occurrence of ripples rather than their overall number.

Given that the interaction between SO-spindles and ripples has been tightly linked to memory reactivation and the behavioural expressions of memory consolidation in rodents ^33,34^, we determined whether TMR-triggered reactivation of head orientation-related activity would be specifically traceable in trials where the probability for SO-spindles and concomitant ripples would be high. Hence, a classifier was trained on the pooled associative retrieval data from both pre- and post-sleep sessions [−0.5 to 1s] and tested on the TMR data [-0.5 to 1.5 s], separately for high SO-spindle activity trials and for low SO-spindle activity trials. The resultant classification performance outcomes were contrasted (see Methods for details). We found a cluster of significant classification from 960 to 1410 ms relative to TMR onset (*p* = 0.019, corrected for multiple comparisons across time, retrieval time-window [-150 to 200 ms]; Fig. 3f; see Supplementary Fig. 9 for results of testing high- and low SO-spindle activity trials against chance-levels and classification results for all TMR segments irrespective of SO-spindle activity). These results indicate that (i) TMR-induced reactivation is related to remembered head orientations and that (ii) reactivation was putatively mediated by SO-spindle and ripple activity. We examine the relation between ripples and memory reactivation in more depth in the next section.

### Spindle-locked MTL ripples facilitate memory reactivation

Having established that cardinal sleep oscillations and reactivation of head orientation-related activity co-occur in time, we next assessed whether ripples and their coupling to spindles would be essential for triggering reactivation processes. First, we tested whether the phase of spindles in cortical contacts would impact ripple band activity in MTL contacts when ripples emerged during the presence of spindles (i.e., 700 to 1400 ms after cue onset; for details see Methods) using the Modulation Index^35^. In line with previous findings, results revealed that the phase of sleep spindles robustly influenced the amplitude in the ripple range^17,21^ (∼ 80 – 120 Hz; *p* = 0.005, corrected for multiple comparisons across frequencies; see Fig. 4a). The phase of cortical delta/theta activity also exhibited a significant effect on ripple activity^36^ (*p* = 0.007, corrected for multiple comparisons across frequencies), while spindle phases additionally modulated low gamma in the MTL (∼20 – 40 Hz; *p* < 0.001, corrected for multiple comparisons across frequencies). When assessing the preferred phase of spindles for their grouping of ripples, we found that ripples were nested towards the trough of cortical spindles (Fig. 4a inset; V-test against ± pi: V = 5.29, *p* = 0.022; mean coupling direction: −176.67 ± 16.61°; mean vector length = 0.21± 0.031)^17,21^.

**Fig. 4.**
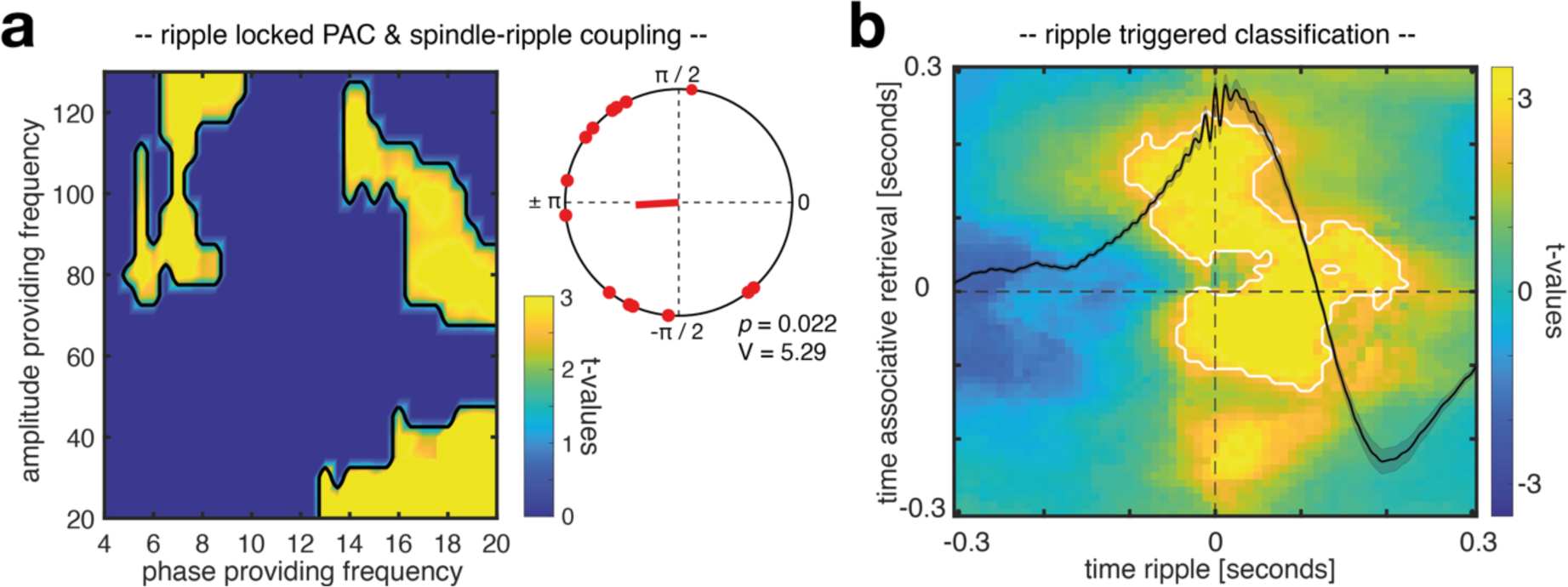
Spindle-ripple interactions and ripple locked classification. (a) Assessing phase-amplitude coupling (PAC) using the Modulation Index revealed that the phase of cortical spindles influenced amplitudes in the ripple range in MTL contacts (∼ 80 – 120 Hz; *p* = 0.005, corrected). In addition, cortical delta/theta phase exhibited a significant effect on MTL ripple amplitudes (*p* = 0.007, corrected), while the spindle phase additionally modulated low gamma amplitudes in the MTL (∼20 – 40 Hz; *p* < 0.001, corrected). The inset illustrates phases of the spindle-ripple modulation, indicating a clustering of ripples towards spindle troughs (corresponding to ±π; V test against ±pi: v = 5.29, p = 0.022; mean coupling direction: −176.67 ± 16.61°, mean vector length = 0.21± 0.031). (b) Head orientation-related brain patterns (left vs. right) were decodable during the presence of spindle locked MTL ripples (contour lines indicate the extent of the significant cluster, *p* = 0.007 corrected; color range (blue to yellow) represents t values).

Finally, we asked whether spindle-locked ripples would be instrumental for memory reactivation to unfold. Hence, again a classifier was trained on the pooled associative retrieval data from both pre- and post-sleep sessions [−0.5 to 1s], but the resulting training weights were this time specifically applied on intracranial EEG segments centered around spindle-locked MTL ripples (i.e., were MTL ripples were paralleled by cortical spindles in between 700 and 1400 ms). For statistical evaluation, surrogate decoding performance was calculated by centering intracranial EEG segments around time-points where no ripple was present during the time-window of preferred spindle-ripple interactions (i.e., 700-1400 ms after cue onset). This procedure was repeated 100 times and resulting surrogate performance values were then averaged, providing baseline values for each participant under the null hypothesis that spindle locked ripples would not be relevant for the classification of stimulus categories. We found a ripple - locked cluster of significant above-chance classification from – 100 to 200 ms relative to ripple centers, indicating that ripples might be indeed facilitating memory reactivation during NREM sleep in humans (*p* = 0.007, corrected for multiple comparisons across time, associative retrieval time-window [-120 to 230 ms], Fig. 4b; see Supplementary Fig. 10 for contrasting ripple triggered classification against chance-level; see Supplementary Fig. 11 for results indicating that uncoupled ripples (i.e., ripples without spindles) did not facilitate multivariate classification).

## Discussion

Our results unveil a key role of spindle-locked ripples in human sleep-based memory reactivation. Specifically, we found that ripples in the MTL, when coupled to cortical spindles, initiate the reprocessing of memories during human NREM sleep, as evidenced by the multivariate classification of prior retrieved head-orientations. These findings elucidate the neural processes mediating memory reactivation during human NREM sleep, by establishing MTL ripples and their synchronization with cortical sleep rhythms as a crucial cornerstone of memory consolidation.

In current models of memory consolidation, ripples are generally considered to be electrophysiological markers of memory reactivation, as they have been suggested to trigger the reprocessing of memories during sleep ^1,20,37^. To date, however, direct evidence for a core contribution of ripples to sleep’s memory function has been lacking in humans. We here used multivariate classification to detect human reactivation processes that are timed by ripples identified in the MTL, providing strong support that ripples in humans initiate memory reactivation akin to animal models ^6,16,38^ and presumably the transfer of memories between the hippocampus and cortical long-term store.

But are all ripples related to memory reactivation? Our data suggest that only a fraction of ripples, specifically those coupled to cortical spindles, were driving the decodability of prior retrieved head-orientations (Fig. 4b and Supplementary Fig. 11). Spindles are well known to group ripples in the MTL ^19,21–23^ (Fig. 4a). They have also been shown to induce neural plasticity in cortical target sites ^13,14,39^, ensuring that those areas are optimally tuned for long-term storage when reactivated memory information arrives ^1^. Hence, our finding that spindle-locked ripples were key for detecting memory reactivation confirms longstanding theoretical predictions concerning the role of synchronized spindle-ripple activity for memory consolidation ^20,37,40^

Moreover, we show that elevated levels of SO-spindle activity promoted the read-out of memory reactivation in both scalp and intracranial EEG recordings. The precise interplay between SOs and spindles is believed to regulate the flow of information between the hippocampus and cortical long-term stores, with SO up-states establishing a time window for spindles and ripples to coincide ^22^. In addition, earlier studies in healthy participants using scalp EEG established SO-spindles as a necessary pre-requisite for the identification of memory reactivation ^18,41,42^. However, because the poorly conducting skull low-pass filters the scalp EEG ^43^, these data remained agnostic to the role of high-frequency signals such as ripples and their potential role in memory reactivation. On basis of our results that ripples in the MTL peaked during the presence of spindles in trials exhibiting high SO-spindle activity (Fig. 3e), while spindle-locked ripples were driving memory reactivation (Fig. 4b), we suggest that also in these previous studies ripples were unbeknownst driving the decodability of prior learned material during SO-spindles.

In the present paradigm, real-world head orientation acted as spatial context in an episodic memory task. By showing that real-world head orientation-related activity is reactivated during successful retrieval and sleep, our findings add ecological validity to prior work on the reactivation of memory contexts ^44–46^. These findings are important because they indicate that the neural correlates of memory functions generalize from screen-based laboratory settings to more naturalistic behavior incorporating bodily movements ^47^. The standard approach to studying the neural basis of human memory requires participants to display minimal bodily movements (e.g., fMRI, MEG), preventing the generation of many self-referential cues, which are thought to play a crucial role in the neural mechanisms underlying memory ^47,48^. The present approach circumvents these shortcomings by incorporating real-world head rotations that trigger self-referential cues such as motor commands, efference copies and reafferent feedback. Combining this approach with rare intracranial recordings from core memory regions (e.g., MTL) opens up exciting opportunities to investigate human electrophysiology that would otherwise remain concealed ^47,49,50^.

On a neural level, little is known about how the human brain tracks and maintains information about real-world head orientation but see ^50^. Animal research, on the other hand, has successfully identified neurons that act as a neural compass during spatial navigation ^51–54^. During sleep, this neural compass seems to be preserved ^55,56^. By simultaneously recording hippocampal ripples and activity from thalamic head direction cells in rodents, Viejo & Peyrache ^57^ showed a specific coupling of the two signals in sleeping rodents that might guide the replay of previously experienced trajectories (even though memory reactivation was not explicitly assessed in their study) ^57^. Our results demonstrating ripple-locked memory reactivation connect to these findings on a conceptual level, by showing that ripples trigger reactivation of memory contexts (i.e., head orientations) that might guide the reactivation of previously experienced events. Going beyond previous work in animal models, we here show that head orientation acts as a memory context in an episodic memory task. Note, however, that the here presented intracranial and surface EEG operate on a meso-/macro scale, compared to the micro scale of single unit recordings. While recent studies have identified a population-level code for real-world head direction ^50,58,59^, future work is necessary to connect the different levels, see e.g. ^60^. On a more general level, implementing real-world navigation into memory paradigms is challenging, but at the same time promises to build bridges between animal research investigating real-world spatial navigation and studies investigating memory processes in humans. Assuming that memory and navigation share neural mechanisms, converging experimental approaches could ultimately foster our understanding of the underlying neural codes in animals and humans ^61^.

We here used TMR as an experimental tool to trigger memory reactivation during sleep. It has been shown that TMR modulates memory leading most often to performance increases ^28,62–66^. Hence, it might seem surprising that TMR did not benefit but deteriorate memory performance both in healthy participants as well as in patients. However, a growing number of studies report TMR-induced impairments, in particular when several targets were associated with a given TMR cue^30,67,68^. In the present study, multiple images (42 in the scalp EEG study; 36 in the intracranial EEG study) were associated with each of the four head orientations. It has been suggested that if the associations between multiple targets and one cue vary in strength, TMR might elicit the reactivation of targets most strongly associated with the cue^67^, akin to models describing retrieval competition during wake ^69,70^. Selectively strengthening a subset of strong cue-target associations via TMR, however, might lead to weakly associated targets losing the competition for being reactivated during a subsequent memory test. Depending on the relative amount of cue-target associations in either subset, this might show as a net beneficial, detrimental, or no effect of TMR on memory performance. Interestingly, in an exploratory analysis (see Supplementary Fig. 12), we found a positive correlation between memory confidence in the pre-sleep memory test and the effect of TMR. Assuming that confidence ratings in our study are positively related to memory strength ^71,72, but see 73^, this relationship indicates that TMR may have mainly reactivated targets that were strongly associated with their cues and further strengthened their association. In turn, these strong targets might have outcompeted weaker ones when competing for being retrieved during post-sleep retrieval, resulting in the observed detrimental effect of TMR on memory performance.

To conclude, using invasive and non-invasive human electrophysiology we found an intimate relationship between NREM sleep related oscillations and memory reactivation. Our findings provide evidence in favor of current models of systems-level consolidation in humans, where spindle-locked ripples synchronize neural population dynamics to reactivate previously formed memories. They establish MTL ripples and their synchronization with cortical sleep rhythms as crucial cornerstones of memory consolidation in humans.

## Methods

### Participants

25 healthy, right-handed participants (mean age: 25.2 ± 0.6; 16 female) with normal or corrected-to-normal vision took part in the EEG experiment. An additional 14 participants had to be excluded due to insufficient sleep or technical problems. The sample size was determined in accordance with previous human sleep and memory studies (e.g., ^74–76^). Pre-study screening questionnaires (including the Pittsburgh Sleep Quality Index (PSQI ^77^), the morningness–eveningness questionnaire ^78^, and a self-developed questionnaire querying general health status and the use of stimulants) indicated that participants did not take any medication at the time of the experimental session and did not suffer from any neurological or psychiatric disorders. All participants reported good overall sleep quality. Furthermore, they had not been on a night shift for at least 8 weeks before the experiment. All participants were instructed to wake up by 7 a.m. and avoid alcohol the evening before and caffeine on the day of the experimental sessions. They received course credit or financial reimbursement in return for their participation. All participants gave written informed consent after being informed about the details of the study. The study was approved by the ethics committee of the Department of Psychology (Ludwig–Maximilian Universität Munich).

For the intracranial EEG study, 10 patients from the Epilepsy Center, Department of Neurology, Ludwig–Maximilian Universität (Munich, Germany), all suffering from medically intractable epilepsy, volunteered (7 female; age: 31.20 ± 3.46). An additional four patients had to be excluded due to technical difficulties. The study was approved by the ethics committee of the Medical Faculty of the Ludwig–Maximilian Universität.

### Stimuli and procedures

#### Overview

On experimental days participants arrived at the sleep laboratory at 7 p.m. The experimental session started with the set-up for polysomnographic recordings during which electrodes for electroencephalographic (EEG) and electrooculography (EOG) were applied. Several days prior to the experimental session, participants were habituated to the environment by having an adaptation nap in the sleep laboratory. At around 8 p.m. the experiment started with a training task, followed by the memory task (for details see Training and Memory Task below). The sleep period began at ∼11 p.m. and all participants slept for ∼7 h (for sleep characteristics see Supplementary Tables 1). During NREM sleep (sleep stages N2 and SWS), some of the animal sounds, which were presented before during the training and the memory task were presented again for 60 minutes (see Targeted memory reactivation for details). Participants were awoken after 6 to 7 hours of sleep from light sleep (sleep stage N1 or N2) and after 15 min of recovery and memory performance was tested again (see Supplementary Table 1 for sleep characteristics).

For the intracranial EEG study the general approach was largely similar. Experimental procedures were arranged around the clinical routines. Specifically, the training sessions were executed during daytime, while the memory task was employed after dinner (i.e., starting between ∼6-7 pm). Patients went to sleep between 10 and 12 p.m. and slept for ∼7 h (for sleep characteristics see Supplementary Tables 2). As with the EEG study, animal sounds were presented for 60 minutes during NREM sleep. Post-sleep memory performance was tested the next morning (see Supplementary Table 2 for sleep characteristics).

#### Stimuli

A set of in total 336 images of objects and five animal sounds (i.e., a cow’s moo, a parrot’s squawk, a cat’s meow, a sheep’s baa, a cuckoo’s sound) served as experimental stimuli. Objects were images of animals, food, clothing, tools, or household items presented on a plain white back-ground. All images were taken from ^79^.

#### Experimental tasks

For the recording of behavioral responses and the presentation of all experimental tasks, Psychophysics Toolbox Version 3 ^80^ and MATLAB 2018b (MathWorks, Natick, USA) were used. Responses were made via keyboard presses on a dedicated PC. Across all experimental phases, presentation order of stimuli was randomized across participants / patients.

#### Training

Participants / patients began by fixating on the center screen, where a fixation cross was presented for 1.5 ± 0.1 seconds. The cross disappeared and one of four animal sounds was played (600 ms). Subsequently, the cross appeared on one of four flanking screens (positioned at −60°, −30°, +30° and 60° relative to the center screen, duration: 2.5 seconds, see Fig. 1a). Four of the total five sounds were randomly chosen at the start of the experiment and randomly assigned to a single flanking screen. The assignment of sound to screen remained fixed across the whole experiment. Participants / patients were instructed to turn their head to face the screen which the fixation cross appeared on and maintain fixation upon the cross (duration: 2.5 seconds). Afterwards, the fixation cross re-appeared on the center screen for 1.5 (± 0.1) seconds and participants had to bring their head back to the starting (i.e., central) position. The training session consisted of 160 trials, split across 4 blocks (i.e., 40 trials per block). The aim of the session was enabling participants / patients to form strong and stable associations between the sound cues and the corresponding head orientations (i.e., flanking screens).

#### Memory task [EEG study]

Participants in the EEG study learned to associate 168 images of objects with specific head orientations. Each trial started with a fixation cross, presented for 1.5 ± 0.1 seconds. Afterwards, one of the four animal sounds from the training phase was played (duration 600ms). Subsequently, an image of an object was presented on the corresponding flanking screen for 4 seconds (the assignment of sound to screen was known to the participants from the training). Participants were instructed to turn their head to face the screen which the image appeared on and to remember the images and their position. Afterwards, the participants had to indicate via button press whether the previously seen object was animate or inanimate, with the question being presented on the center screen. The pre-sleep memory test included the 168 images from encoding (old items) intermixed with 84 new images, which were not seen by the participants before (“foils”). Each trial started with a fixation cross, presented for 1.5 ± 0.1 s. After the fixation cross, an image was presented on the center screen. After 1 second, participants were asked to indicate whether the image was “old” (i.e., part of the learning material) or “new”’ (i.e., it was not seen during learning) within the next 10 s. In case of “new” responses, participants immediately moved on to the next trial. In case of “old” responses, participants were required to indicate by button press the related head orientation (i.e., the flanking screen on which the image was presented). Each trial ended with participants indicating how confident they were with their head orientation decision (scale from 0 corresponding to very uncertain to 4, very certain). The post-sleep retrieval followed the same procedures as the pre-sleep memory test with the exception that new foil images were used.

#### Memory task [intracranial EEG study]

The procedures of the memory task were similar to the EEG study, with some modifications. The stimulus pool was comprised of 288 objects (drawn from the same selection as used in the EEG study). In order not to overtax patients, the pre-sleep memory task was split into three consecutive encoding - retrieval blocks. During each encoding block patients were presented with 48 images on the flanking screens (please note that the trial level was identical to the one described above). Each encoding block was followed by a retrieval block, where the 48 images from encoding intermixed with 24 new images were presented. Hence, across all blocks, we used 144 images as old items and 72 images as new items. As above, patients had to first indicate whether a given image was old or new and in case of old items specify the remembered head orientation. Due to time constraints no confidence rating was obtained. The post-sleep retrieval was executed in one run, meaning that patients were confronted with the 144 images which were part of the learning material and 72 foils.

#### Targeted memory reactivation

For targeted memory reactivation 2 out of the 4 sounds presented during training and encoding were selected. Specifically, we randomly picked one out of the two sounds associated with the left-sided head orientations (i.e., flanking screens positioned at −60° and −30°) and one sound associated with the right-sided head orientations (i.e., flanking screens positioned at 30° and 60°). In addition, the fifth animal sound which was not used during training and encoding served as a control stimulus. The three cues were repeatedly presented during NREM sleep via loudspeaker with an intertrial interval of 5.5 ± 0.2 seconds (∼50 dB sound pressure level) for a maximum of 60 minutes (EEG study: 182.6 ± 31.41 repetitions per stimulus; intracranial EEG study: 187.1 ± 23.7 repetitions per stimulus). Sound presentation was stopped whenever signs of arousals, awakenings or REM sleep became visible.

##### Scalp EEG acquisition

An EEGo 65 channel EEG system (ANT Neuro Enschede, Netherlands) was used to record electro-encephalography (EEG) throughout the experiment. Impedances were kept below 20 kΩ. EEG signals were referenced online to electrode CPz and sampled at a rate of 1000 Hz. Furthermore, horizontal and vertical EOG was recorded for polysomnography. Sleep architecture was determined offline according to standard criteria by two independent raters ^81^.

##### intracranial EEG acquisition

Intracranial EEG was recorded from Spencer depth electrodes (Ad-Tech Medical Instrument, Racine, Wisconsin, United States) with 4–12 contacts each, 5 mm apart. Data were recorded using XLTEK Neuroworks software (Natus Medical, San Carlos, California, US) and an XLTEK EMU128FS amplifier, with voltages referenced to a parietal electrode site. The sampling rate was set at 1000 Hz.

##### EEG data analysis

EEG data were preprocessed using the FieldTrip toolbox for EEG/MEG analysis ^82^. All data were down-sampled to 200 Hz. Subsequently, the pre- and post-sleep retrieval as well as the TMR data were segmented into epochs. For the retrieval data, we segmented data from the onset of the associative retrieval. We reasoned that memory reactivation of associated head orientation-s should be particularly strong due to the potential hippocampal dependency (as compared to recognition tests ^83^).. The temporal range of the epochs was [–1 to 3 s] around stimulus onset for retrieval and TMR trials. Noisy EEG channels were identified by visual inspection, discarded, and interpolated, using a weighted average of the neighboring channels. The data were visually inspected and artefactual trials were removed. The retrieval data were additionally subjected to an independent component analysis (ICA) and ICA components associated with eye blinks and eye movements were identified and rejected.

##### intracranial EEG data analysis

The preprocessing steps for the intracranial EEG data were identical to the ones described above, just that intracranial EEG data were additionally inspected for epileptic activity, with data segments comprising epileptic events at any given contact being discarded (36.14 ± 5.51 % of all trials (N_allTrials_ = 633 ± 26.48); for interictal epileptiform discharge triggered classification see Supplementary Fig. 13). In addition, contacts which were contaminated with epileptiform background activity were discarded. Only seizure-free nights were included in the analysis.

##### Source level

To estimate the sources of the obtained effects in the scalp EEG study, we applied a DICS beamforming method ^84^, as implemented in FieldTrip ^82^. A spatial filter for each specified location (each grid point; 10mm^3^ grid) was computed based on the cross-spectral density, calculated separately for all retrieval and TMR trials. Electrode locations for the 65-channel EEGo EEG system were co-registered to the surface of a standard MRI template in MNI (Montreal Neurological Institute) space using the nasion and the left and right preauricular as fiducial landmarks. A standard leadfield was computed using the standard boundary element model ^39^. The forward model was created using a common dipole grid (10mm3 grid) of the grey matter volume (derived from the anatomical automatic labeling atlas ^85^ in MNI space, warped onto standard MRI template, leading to 1457 virtual sensors. Data analysis was accomplished in the same way as on sensor level.

##### Time–frequency analysis

Time–frequency analysis of the TMR segments (memory related and control cues) was performed using FieldTrip. Frequency decomposition of the data, using Fourier analysis based on sliding time windows (moving forward in 50 ms increments). The window length was set to five cycles of a given frequency (frequency range: 1–25 Hz in 1 Hz steps). The windowed data segments were multiplied with a Hanning taper before Fourier analysis. Afterwards, power values were z-scored across time [−1 to 3 s]. The longer time segments were chosen to allow for resolving low frequency activity within the time windows of interest [−0.5 to 1.5 s] and avoid edge artifacts. For intracranial EEG data frontal, parietal and temporal contacts were taken into account.

##### Multivariate analysis

Multivariate classification of single-trial EEG data was performed using MVPA-Light, a MATLAB-based toolbox for multivariate pattern analysis ^86^. For all multivariate analyses, a LDA was used as a classifier. Prior to classification, data in both studies were re-referenced using a common average reference (CAR).

For classification within the retrieval task, the localizer data were z-scored across all trials for each time point separately. Next, data from the pre- and the post-sleep retrieval were collapsed and subjected to a principal component analysis (PCA), which transforms the data into linearly uncorrelated components, ordered by the amount of variance explained by each component ^87^. PCA was applied to reduce dimensionality and limit over-fitting (PCA) and the first 30 principal components were retained for further analysis. To quantify whether remembered head orientations can be differentiated during retrieval, the classifier was trained and tested to discriminate between the later cued head orientations (i.e., one left sided and one right sided head orientation; see Targeted Memory reactivation for details). Only trials belonging to remembered head orientations entered the analysis. Data were smoothed using a running average window of 150 ms. The EEG channels / intracranial EEG contacts served as features and a different classifier was trained and tested on every time point. As metric, we used Area Under the ROC Curve (AUC), which indexes the mean accuracy with which a randomly chosen pair of Class A and Class B trials could be assigned to their correct classes (0.5 = random performance; 1.0 = perfect performance). To avoid overfitting, data were split into training and test sets using fivefold cross-validation ^88^. Since cross-validation results are stochastic due to the random assignment of trials into folds, the analysis was repeated five times and results were averaged. For statistical evaluation, the classification output was tested against chance levels (i.e., 0.5). To resolve the topography of diagnostic features in the scalp EEG data, we conducted a “searchlight decoding procedure” (Fig. 2c). In brief, PCA components were projected back to sensor space and the classification procedure was repeated across moving kernels of small electrode clusters, with neighboring electrodes being selected as features [feature number range: 5 to 9]. Finally, classification values were collapsed across our time windows of interest [retrieval time: −30 to 680 ms;] and tested against chance level (corrected for multiple comparisons across space).

To investigate whether TMR would elicit head orientation-related activity, we used the temporal generalization method ^89^. Prior to decoding, a baseline correction was applied based on the whole trial for retrieval and TMR segments [−0.5 to 3 s]. Next, retrieval and TMR data were z-scored across trials and collapsed. PCA was applied to the pooled retrieval-TMR data and the first 30 principal components were retained. Retrieval and TMR data were smoothed using a running average window of 150 ms. A classifier was then trained for every time point in the retrieval data and applied on every time point during TMR. No cross-validation was required since retrieval and TMR datasets were independent. As metric, we again used AUC (see above). For statistical evaluation, the classification output was tested against chance levels (i.e., 0.5).

Given that the interaction between SO-spindles and ripples has been tightly linked to memory reactivation^33,34^, we determined whether TMR-triggered reactivation of head orientation activity would be traceable in intracranial EEG recordings, specifically in trials where the probability for SO-spindles and concomitant ripples would be high: for each participant, we sorted the TMR trials as a function of power in the clusters obtained in the time-frequency analysis (Fig. 3c) and divided the trials using a median split. Then, a classifier was trained on the concatenated retrieval data from both pre- and post-sleep sessions [−0.5 to 1s] and the resulting training weights were applied on the TMR data [-0.5 to 1.5 s], either comprising high SO-spindle power trials (i.e., where ripples peaked during spindle activity) or low SO-spindle power trials and contrasted the resultant performance outcomes. For statistical evaluation, classification performance of both categories was directly compared.

For ripple triggered classification, a classifier was trained on the concatenated retrieval data from both pre- and post-sleep sessions [−0.5 to 1s], but the resulting training weights were applied on intracranial EEG segments centered around spindle-locked MTL ripples (i.e., ripples occurring during spindles between 700 to 1400 ms after cue onset). For statistical evaluation, surrogate decoding performance was calculated by centering intracranial EEG segments around time-points where no ripple was present during the time-window of preferred spindle-ripple interactions (i.e., 700-1400 ms after cue onset). This procedure was repeated 100 times and resulting surrogate performance values were then averaged, providing baseline values for each participant under the null hypothesis that spindle locked ripples would not be relevant for the classification of stimulus categories.

For the scalp EEG data, head orientations were decoded in source space using searchlight analysis ^90^. A sphere of radius 2 cm was centered on each of the 1467 voxels in the brain. All voxels within the sphere that were inside the brain volume (10-26 voxels) were selected as features. Identical to the sensor level analysis a classifier was trained for every time point in the retrieval data and applied on every time point during TMR. Finally, classification values were collapsed across our time windows of interest [retrieval time: −110 to 330 ms;] and tested against chance level (corrected for multiple comparisons across space).

##### Ripple detection

Ripple events in the medial temporal lobe (MTL) depth recordings (7 patients, 14 contacts in total: 4 hippocampal, 7 parahippocampal and 3 entorhinal contacts) were detected during artifact-free TMR segments using offline algorithms ^18,91^. The intracranial EEG signal (sampling rate 1000 Hz) was band-pass filtered from 80 to 120 Hz and the root mean square signal (RMS) was calculated based on a 20 ms windows followed by an additional smoothing with the same window length. A ripple event was identified whenever the smoothed RMS-signal exceed a threshold, defined by the mean plus 2 times the standard deviation of the RMS-signal across all TMR data points. Potential ripple events shorter than 25 ms or longer than 300 ms were rejected. All ripple events were required to exhibit a minimum of three cycles in the raw signal.

##### Peri-event histograms of ripple occurrence

To investigate the timing of MTL ripples (centered at the maximal negative amplitude) with regards to TMR cues, we first sorted for each participant the TMR trials as a function of power in the clusters obtained in the time-frequency analysis (Fig. 3c) and divided the trials using a median split. We then created for each condition peri-event histograms (bin size = 50 ms) of ripple events time-locked to TMR cues. The resulting histograms were normalized by the total number of TMR trials (multiplied by 100).

##### Modulation Index

Phase amplitude coupling was assessed with the Modulation Index (MI)^35^. We first isolated in each patient the cortical contact exhibiting the strongest power in the spindle band (12-15 Hz; 0 – 1.5 seconds after cues onset; see Supplementary Table 3 for an overview). All intracranial EEG data segments were centered in relation to MTL detected ripple maxima, focusing on ripples which paralleled cortical spindles (i.e., ripples emerging in a time-window from 700 to 1400 ms after cue onset). Low frequencies in cortical contacts (4 – 20 Hz) were filtered with a window of 0.3 times the frequency of interest, centered on each frequency step. High frequencies in MTL contacts (20 – 130 Hz) were filtered with a window of 0.7 times the frequency of interest. To compute the MI (for a given frequency pair), we divided the phase signal into 18 bins (20° each), and then, computed for each bin the mean amplitude. This yielded a distribution of amplitude as a function of phase. The MI is defined as the Kullback-Leibler distance between that distribution and the uniform distribution (over the same number of bins). To assess the statistical significance of the MI values, we randomly shuffled the trials of the amplitude providing contacts and computed the MI using the shuffled data. We repeated this procedure 100 times, resulting in a MI-level reference distribution.

##### Spindle-ripple coupling

For the analysis of the coupling between cortical spindles and MTL ripples, we first isolated in each patient the cortical contact exhibiting the strongest power in the spindle band (12-15 Hz; 0 – 1.5 seconds after cues onset). We then filtered the data (12-15 Hz, two-pass Butterworth bandpass filter) and applied a Hilbert transform. The instantaneous phase angle of cortical recordings at the time of MTL detected ripple maxima was extracted. We specifically focused on ripples which occurred during cortical spindles (i.e., ripples emerging in a time-window from 700 to 1400 ms after cue onset). The preferred phase of spindle-ripple coupling for each cortical contact was then obtained by taking the circular mean of all individual events’ preferred phases.

##### Statistics

Behavioral retrieval data were subjected to a 2 (TMR: cued/uncued) × 2 (Test-Time: Pre-sleep/Post-sleep) repeated measures ANOVA. The statistical significance thresholds for all behavioral analyses were set at *p* < .05. FieldTrip’s cluster permutation test ^82^ was used to deal with the multiple comparisons problem for all classification analyses. A dependent-samples t-test was used at the sample level to identify clusters of contiguous time points across participants and values were thresholded at *p* = 0.05. Monte Carlo simulations were used to calculate the cluster p value (alpha = 0.05, two-tailed) under the permutation distribution. Analyses were performed at the group level. The input data were either classification values across time (Fig. 1c + 3b) or time x time classification values (Fig. 2b + 3c). In all cases a two-sided cluster permutation test with 1000 randomizations was used to contrast classification accuracy against chance performance. The same statistical rationale was implemented for the statistical assessment of time frequency data with time × frequency values as input, as well as for phase-amplitude data (frequency x frequency as input) and peri-event histograms (time as input). Statistical analysis of TFR data in the intracranial EEG study was performed at the individual electrode/contact level (fixed-effects analysis), considering all intracranial EEG contacts (N = 389; see Supplementary Fig. 9 for coverage), while statistical analysis of phase-amplitude and peri-event histogram data considered all possible cortical-MTL contact pairs (N = 14, fixed effects; with chosen cortical contacts showing the strongest spindle power). Pearson correlation was used to assess the relationship between (i) classification performance and time-frequency power (Fig. 2d) and (ii) the time-course of classification performance and ripple density (Fig. 3d). For circular statistics (Fig. 4a), the phase distributions across all cortical-MTL contact pairs (N = 14) were tested against uniformity with a specified mean direction (i.e., ±π corresponding to the spindle through) using the V-test (CircStat toolbox^92^).

## Acknowledgements

This work was supported by the European Research Council (https://erc.europa.eu/, Starting Grant 802681 awarded to T.St). T.S is supported by the Emmy Noether program of the German Research Foundation (492835154). We are indebted to all patients and participants who volunteered to participate in this study. We thank the staff and physicians at the Epilepsy Center, Department of Neurology, Ludwig Maximilians University, Munich for assistance.

## Competing interests

The authors declare no competing interests.

## Supplementary Information

**Supplementary Figure 1.**
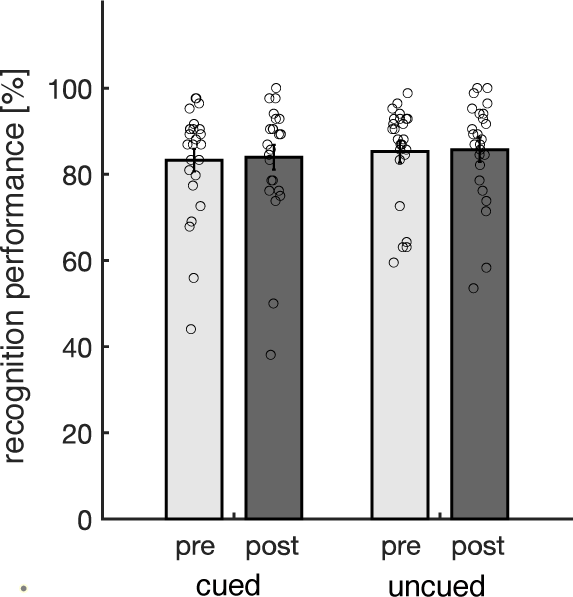
Recognition memory performance (EEG study). Behavioral results for both experimental sessions pre- (light gray) and post-sleep (dark gray), separated into cued and uncued trials. Bar graphs show mean (±SEM) percentage of correctly recognized images (‘hits’). Dots indicate individual memory performance of participants (N = 25). There was neither a significant main effect of test time (F1,24 = 0.29; *p* = 0.59), nor a significant interaction between test-time and cueing (F1,24 = 0.08; *p* = 0.77).

**Supplementary Figure 2.**
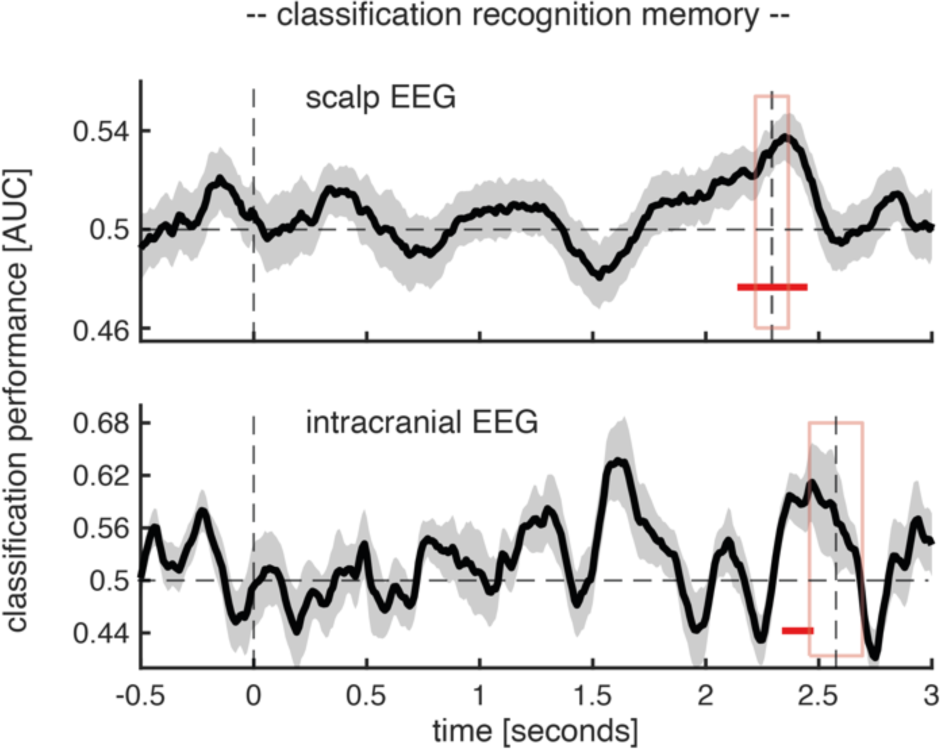
Classification of later cued head orientations (left vs. right) locked to recognition onset (time = 0; first dashed vertical line). Later cued head orientations could be decoded (above chance) towards the end of recognition test trials, briefly preceding the onset of the of the associate prompt. The second vertical dashed line indicates the mean onset of the associative memory prompt. The red rectangle illustrates the standard error of the mean. The black solid line indicates decoding performance (±SEM). The horizontal dashed line indicates chance level performance (i.e., 0.5). The lower horizontal red line shows the temporal extent of significant decoding results as derived from a dependent-samples t-test (one-sided, scalp EEG study: p < 0.032; intracranial EEG study: p = 0.043, cluster corrected across time).

**Supplementary Figure 3.**
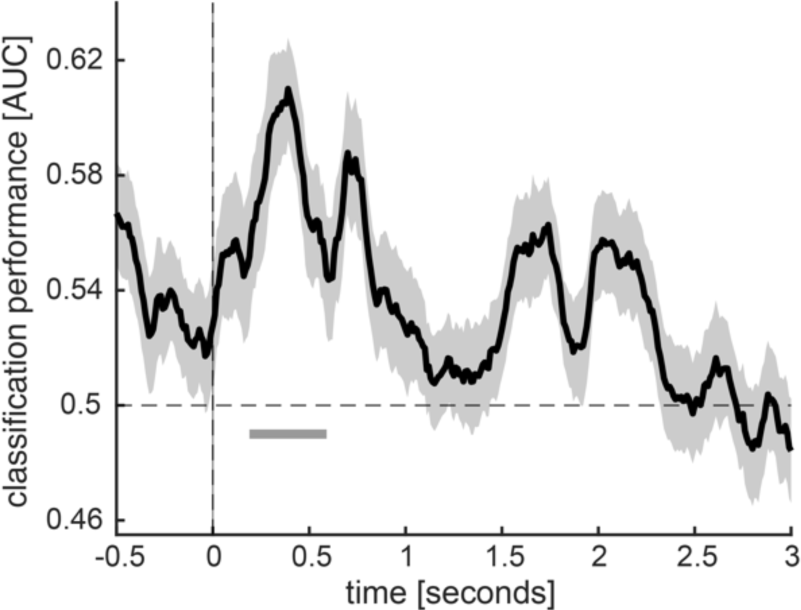
Classification of later not cued head orientations during retrieval. Later not cued head orientations (left vs. right) could be reliably decoded (above chance) from the retrieval data between 190 and 590 ms after the onset of the associate prompt (the black solid line indicates decoding performance (±SEM)). The horizontal dashed line indicates chance level performance (i.e., 0.5). The vertical solid line indicates the onset of associative retrieval trials (time = 0). The lower horizontal gray line shows the temporal extent of significant decoding results as derived from a dependent-samples t-test (two-sided, *p* < 0.006, cluster corrected across time).

**Supplementary Figure 4.**
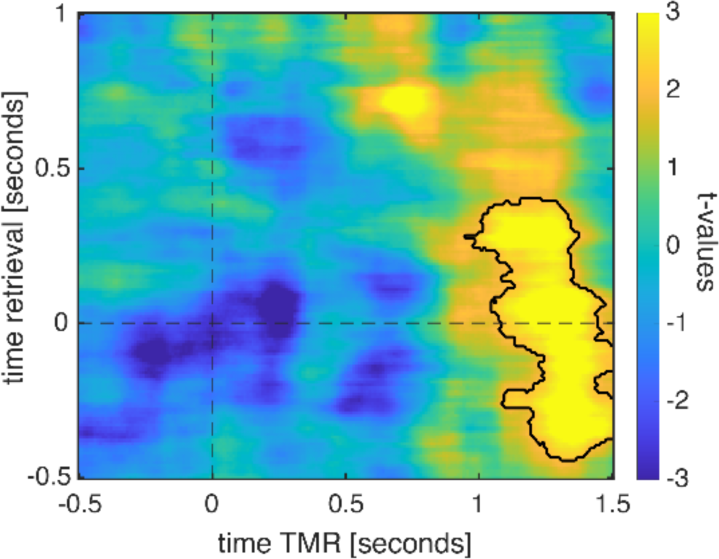
Classification using TMR trials exhibiting increased levels of activity in the SO-spindle range. Retrieval-related brain patterns (left vs. right head orientations) were reliably decodable during ‘high power’ TMR trials (contour lines indicate the extent of the significant cluster, *p* = 0.005 corrected; color range (blue to yellow) represents t values against chance level performance (i.e., 0.5)).

**Supplementary Figure 5.**
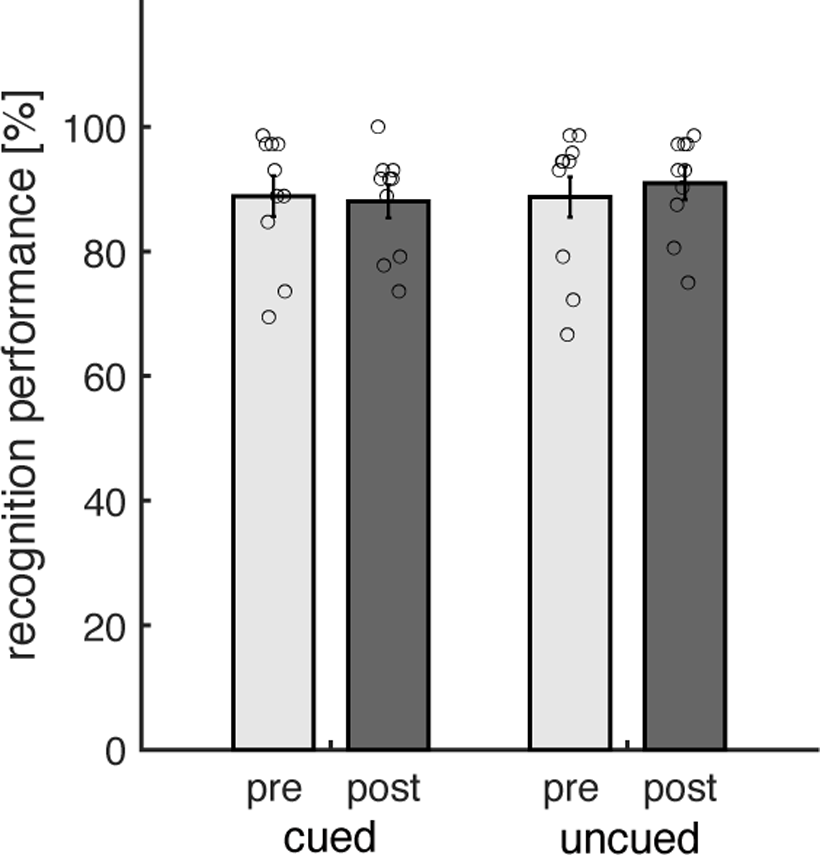
Recognition memory performance (intracranial EEG study). Behavioral results for both experimental sessions pre- (light gray) and post-sleep (dark gray), separated into cued and uncued trials. Bar graphs show mean (±SEM) percentage of correctly recognized images (‘hits’). Dots indicate individual memory performance of participants (N = 10). There was neither a significant main effect of test time (F1,9 = 0.06; *p* = 0.08); nor a significant interaction between test-time and cueing (F1,9 = 2.25; *p* = 0.16).

**Supplementary Figure 6.**
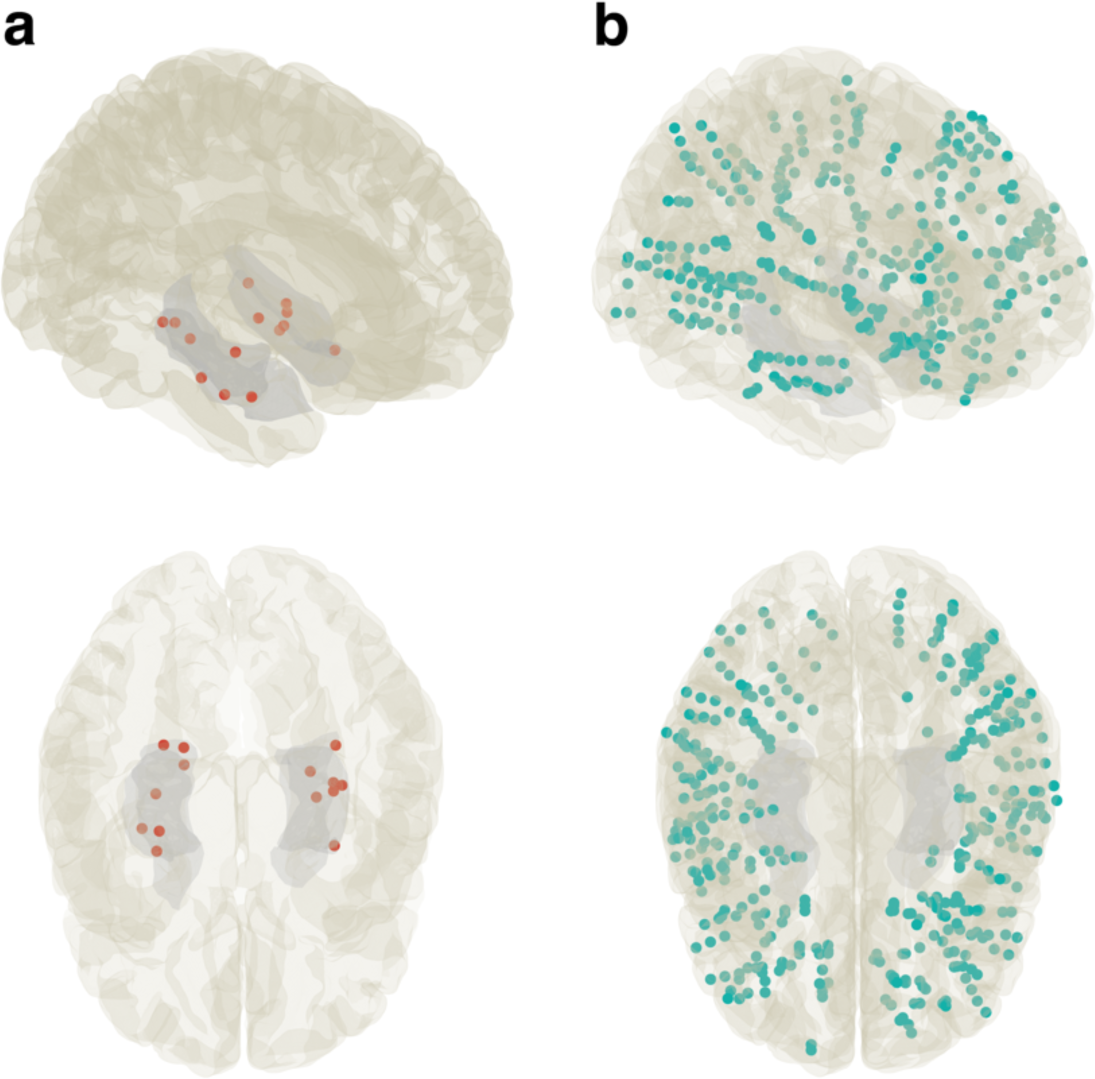
(a) Group-level (N = 7) electrode coverage in MNI space of intracranial electrodes in the MTL (comprising contacts in the hippocampus, parahippocampus and entorhinal cortex; N = 14; red). Group-level (N = 10) electrode coverage in MNI space of intracranial electrodes in non-MTL cortical areas (N = 14; 157 frontal; 55 parietal, 121 temporal, 42 occipital contacts).

**Supplementary Figure 7.**
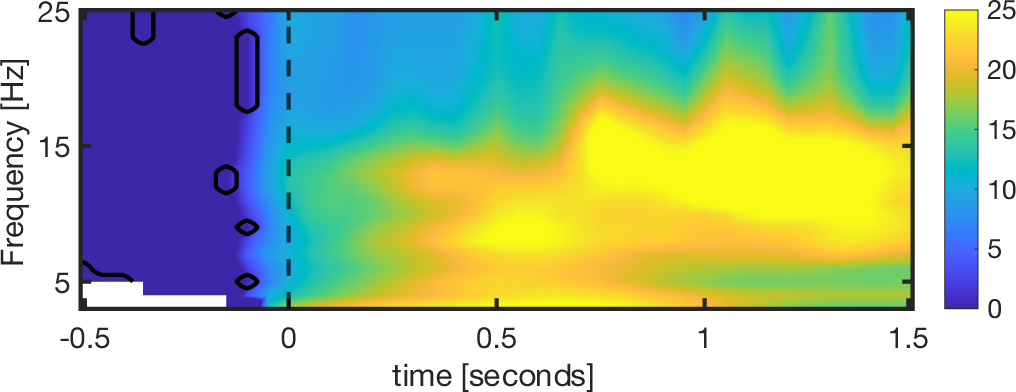
Time–frequency representation difference map of high and low SO-spindle activity trials. Oscillatory power was significantly higher for all time and frequency bins starting around cue onset for high power trials. Note the low frequency activity around 500 ms and subsequent spindle activity standing out (12-16Hz; *p* < 0.00001, corrected for multiple comparisons across time and frequency).

**Supplementary Figure 8.**
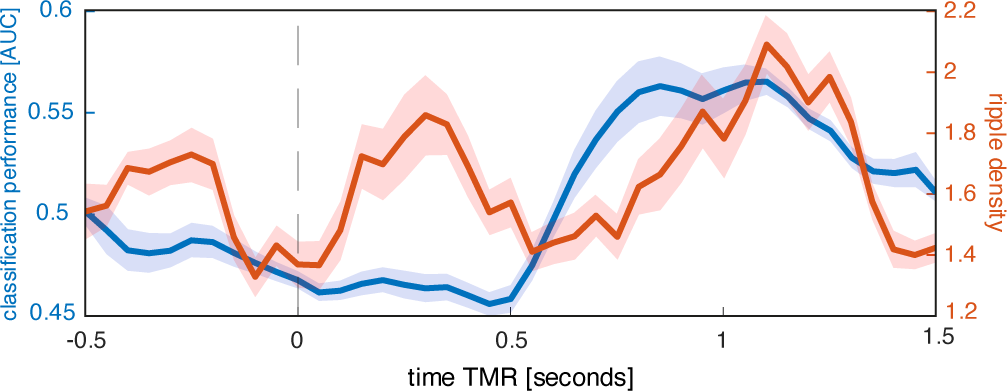
Time course of ripple density and reactivation signal. Blue: Classification output averaged across the relevant retrieval time [-150 to 200 ms]. Red: Patient-averaged ripple density across time.

**Supplementary Figure 9.**
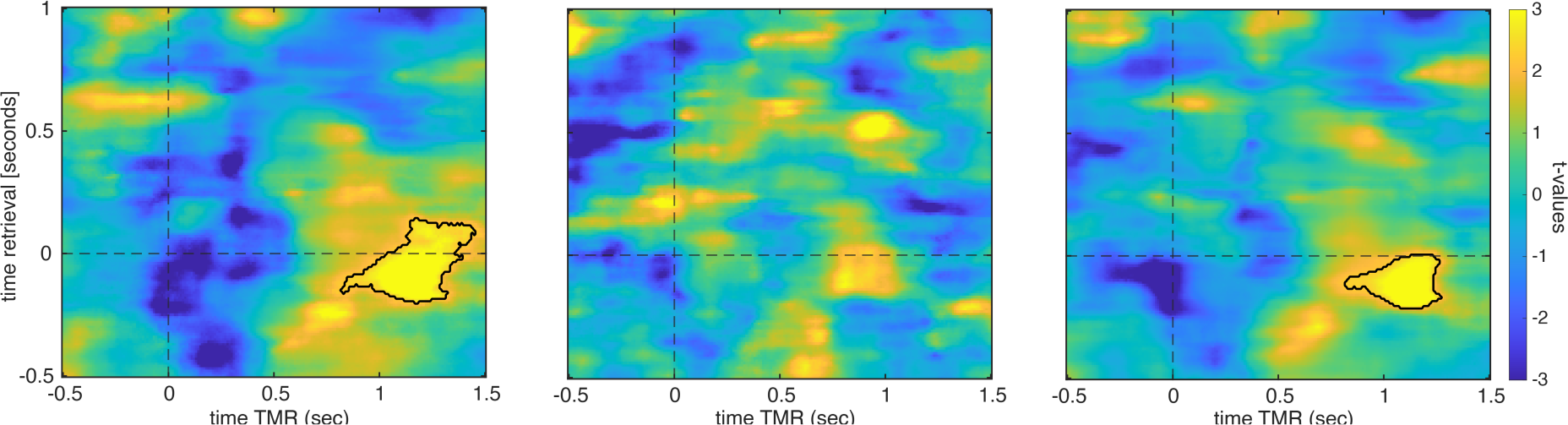
(left) Classification using TMR trials exhibiting increased levels of activity in the SO-spindle range. Retrieval-related brain patterns (left vs. right head orientations) were reliably decodable during ‘high power’ TMR trials (contour lines indicate the extent of the significant cluster, p = 0.004 corrected; color range (blue to yellow) represents t values against chance level performance (i.e., 0.5)). (middle) Classification using TMR trials exhibiting low levels of activity in the SO-spindle range. Retrieval-related brain patterns (left vs. right head orientations) were not decodable during ‘low power’ TMR trials (contour lines indicate the extent of the significant cluster, p = 0.71 corrected; color range (blue to yellow) represents t values against chance level performance (i.e., 0.5)). (right) Classification using all TMR trials (irrespective of SO-spindle power). Retrieval-related brain patterns (left vs. right head orientations) were reliably decodable during ‘high power’ TMR trials (contour lines indicate the extent of the significant cluster, *p* = 0.047 corrected; color range (blue to yellow) represents t values against chance level performance (i.e., 0.5)).

**Supplementary Figure 10.**
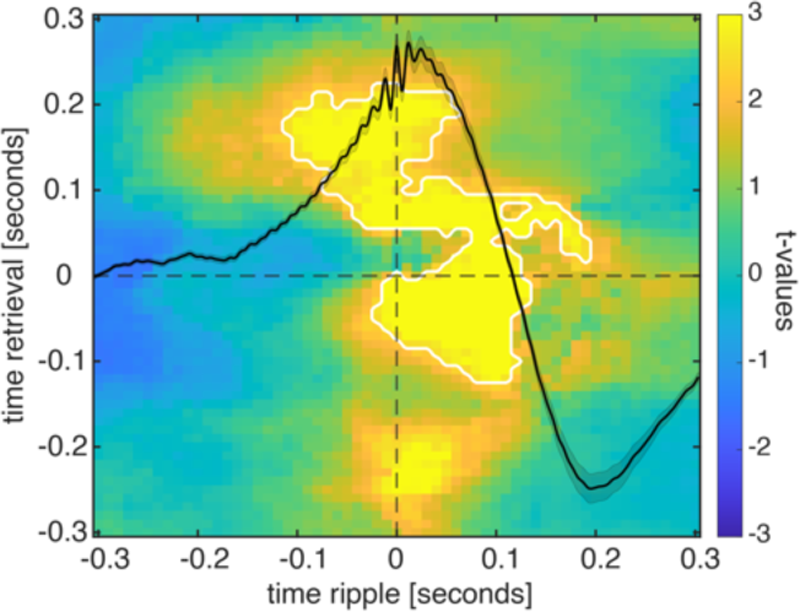
Retrieval-related brain patterns (left vs. right head orientations) were decodable during the presence of spindle locked MTL ripple when tested against chance level (i.e., 0.5; contour lines indicate the extent of the significant cluster, *p* = 0.007 corrected; color range (blue to yellow) represents t values).

**Supplementary Figure 11.**
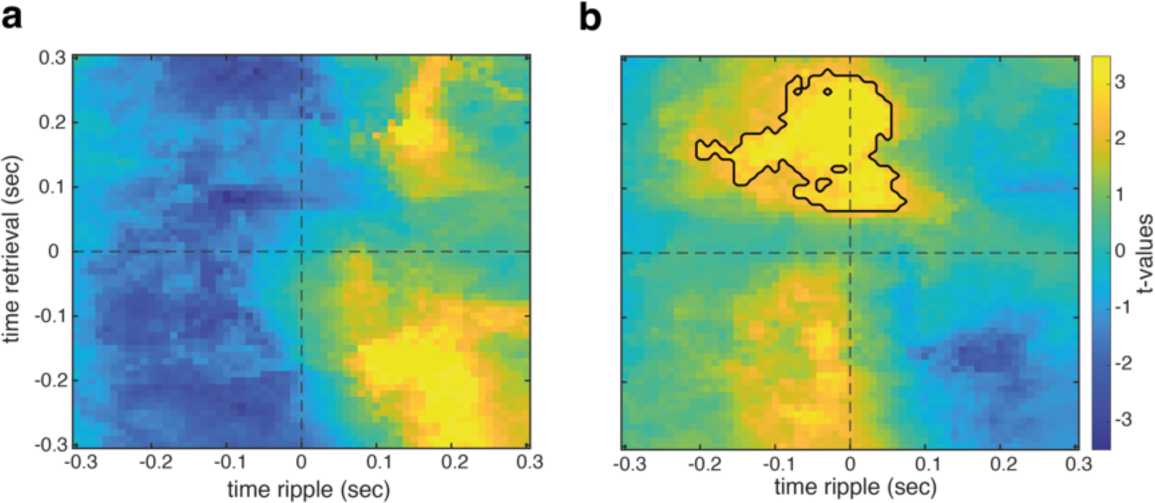
(a) Retrieval-related brain patterns brain patterns (left vs. right head orientations) were not decodable during the presence of uncoupled ripples (i.e., ripples without spindles; *p* = 0.073). (b) Head orientation-related brain patterns (left vs. right) were decodable during TMR when contrasting data segments centered on spindle-locked ripples and uncoupled ripples (contour lines indicate the extent of the cluster, p = 0.032 corrected; color range (blue to yellow) represents t values against chance level performance).

**Supplementary Figure 12.**
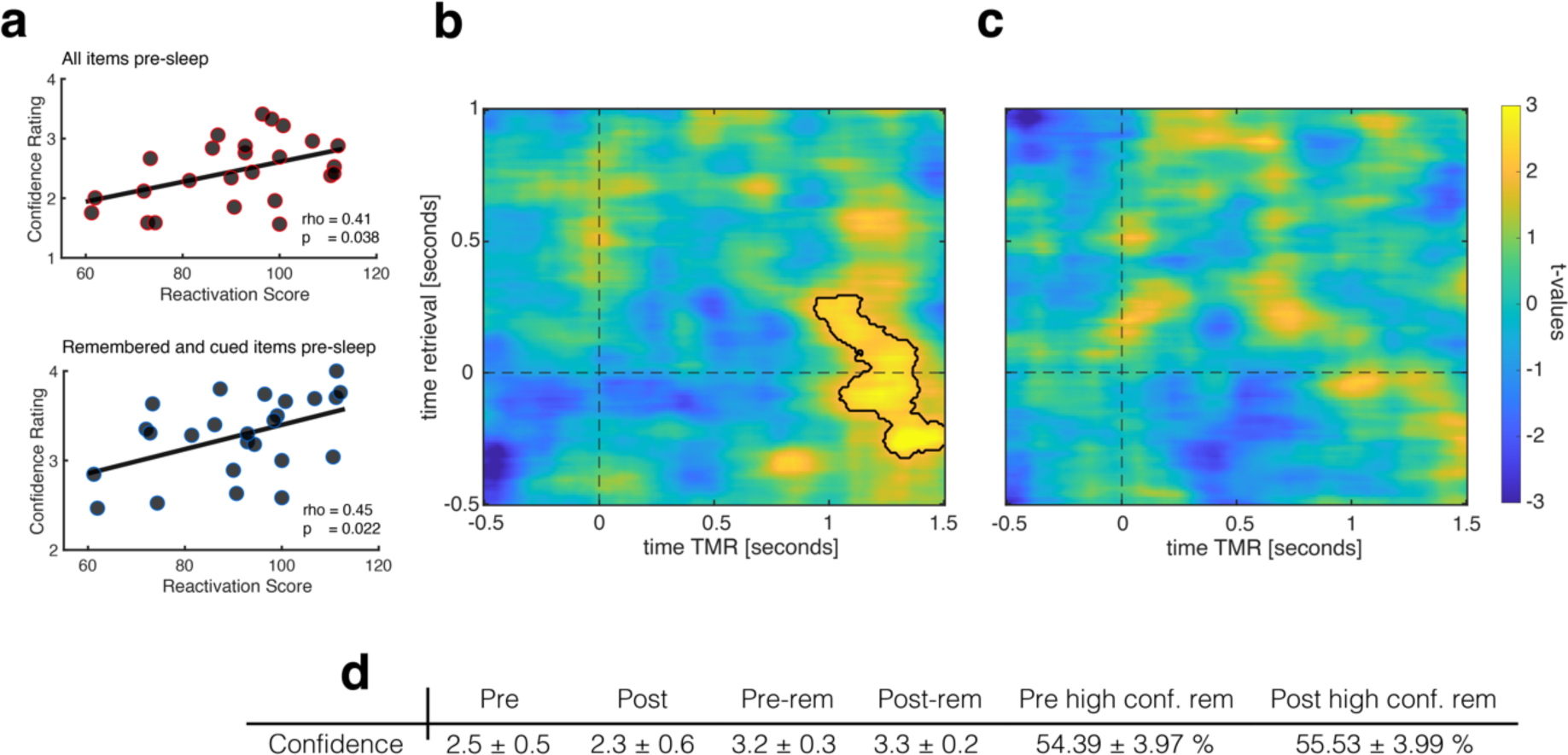
Relationship of confidence ratings and memory reactivation. In each retrieval trial, participants indicated how confident they were with their head orientation decision (scale from 0 corresponding to very uncertain to 4, very certain). (a) Participants’ average confidence ratings across all trials correlated positively with the behavioral impact of TMR (i.e., reactivation score: relative difference from pre- to post-sleep for cued items - relative difference from pre- to post-sleep for uncued items * 100)/ pre-sleep remembered items; rho = 0.41; *p* = 0.038). Likewise, participants’ average confidence ratings across all remembered and cued trial correlated positively with the behavioral impact of TMR (i.e., reactivation score: relative difference from pre- to post-sleep for cued items - relative difference from pre- to post-sleep for uncued items * 100)/ pre-sleep remembered items; rho = 0.41; *p* = 0.038). These results suggest that highly confident participants rather benefitted from TMR, while less confident participants exhibited a detrimental effect of TMR. (b) Classification using only high confidence trials (confidence rating = 4) as training data. Retrieval-related brain patterns (left vs. right head orientations) were reliably decodable when the decoder was trained on high confidence trials (contour lines indicate the extent of the significant cluster, *p* = 0.025 corrected; color range (blue to yellow) represents t values against chance level performance (i.e., 0.5)). (c) Classification using lower confidence trials (confidence rating < 4) as training data. Retrieval-related brain patterns (left vs. right head orientations) were not decodable when the decoder was trained on lower confidence trials (*p* = 0.96 corrected; color range (blue to yellow) represents t values against chance level performance (i.e., 0.5)). (d) Descriptives of confidence ratings: Means and SEM of confidence ratings of all pre-sleep trials [Pre], all post sleep trials [Post], all remembered pre-sleep trials [Pre-rem] and all remembered post sleep trials [Post-rem]. Percentage of high confidence trials of all remembered trials in the pre-sleep memory test [Pre high conf. rem] and percentage of high confidence trials of all remembered trials in the post-sleep memory test [Post high conf. rem].

**Supplementary Figure 13.**
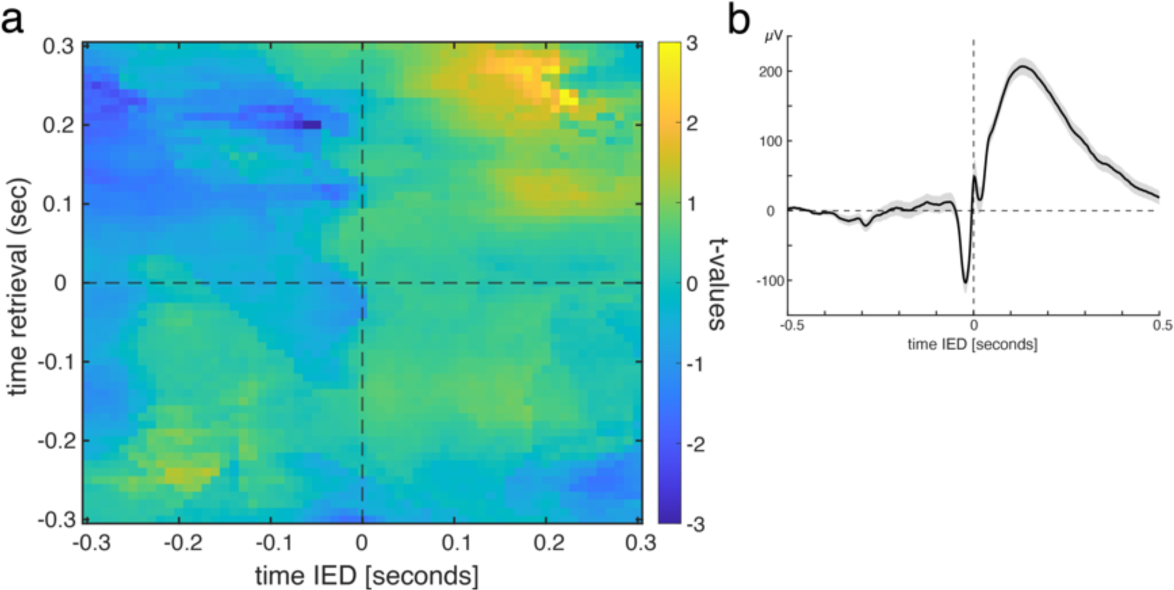
Interictal epileptiform discharge (IED) triggered classification. IEDs were automatically detected in MTL contacts of discarded data segments using established algorithms ^17^. All intracranial EEG segments were centered around IEDs emerging 700 to 1400 ms after stimulus onset (i.e., same convention as in the ripple-triggered classification). (a) Training a classifier on the pooled associative retrieval data from both pre- and post-sleep sessions [−0.5 to 1s] and testing on the IED centered intracranial EEG data did not yield any significant result when tested against chance level performance (i.e., 0.5; *p* = 0.71). (b) Single subject example of IED centered ERP.

**Supplementary Table 1.** Sleep characteristics EEG study. Data are means ± s.e.m. N1, N2: NREM sleep stages N1 & N2, SWS: slow-wave sleep, REM: rapid eye movement sleep, WASO: wake after sleep onset.

|  | N1 | N2 | SWS | REM | WASO | TST [min] |
| --- | --- | --- | --- | --- | --- | --- |
| Sleep stage [%] | 4.1 ± 0.5 | 46.8 ± 1.4 | 22.6 ± 1.0 | 21.0 ± 1.2 | 0.04 ± 0.01 | 421.4 ± 9.7 |

**Supplementary Table 2.** Sleep characteristics iEEG study. Data are means ± s.e.m. N1, N2: NREM sleep stages N1 & N2, SWS: slow-wave sleep, REM: rapid eye movement sleep, WASO: wake after sleep onset.

|  | N1 | N2 | SWS | REM | WASO | TST [min] |
| --- | --- | --- | --- | --- | --- | --- |
| Sleep stage [%] | 4.1 ± 0.9 | 44.9 ± 2.7 | 20.6 ± 2.5 | 23.1 ± 2.0 | 5.2 ± 1.7 | 480.3 ± 31.4 |

**Supplementary Table 3.**
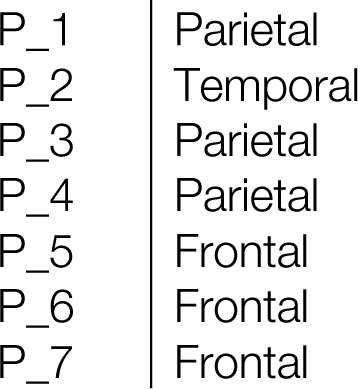
Location of cortical contact exhibiting the strongest power in the spindle band (12-15 Hz)

## Notes

### Competing Interest Statement

The authors have declared no competing interest.

### Summary of Updates

- Results section 'Spindle-locked MTL ripples facilitate memory reactivation' and Fig. 4 added to assess the specific role of ripples in memory reactivation. - Supplemental Figures updated

